# Alteration of the premature tRNA landscape by gammaherpesvirus infection

**DOI:** 10.1101/2019.12.22.886382

**Authors:** Jessica Tucker, Aaron M. Schaller, Ian Willis, Britt A. Glaunsinger

**Affiliations:** Department of Plant and Microbial Biology, University of California Berkeley, CA, USA; Department of Molecular and Cell Biology, University of California Berkeley, CA, USA; Departments of Biochemistry and Systems and Computational Biology, Albert Einstein College of Medicine, Bronx NY, USA; Howard Hughes Medical Institute, University of California Berkeley, CA, USA

## Abstract

Transfer RNAs (tRNAs) are transcribed by RNA polymerase III (RNAPIII) and play a central role in decoding our genome, yet their expression and non-canonical function remain understudied. Many DNA tumor viruses enhance the activity of RNAPIII, yet whether infection alters tRNA expression is largely unknown. Here, we present the first genome-wide analysis of how viral infection alters the tRNAome. Using a tRNA-specific sequencing method (DM-tRNA-seq), we find that the murine gammaherpesvirus MHV68 induces global changes in pre-tRNA expression with 14% of tRNA genes upregulated more than 3-fold, indicating that differential tRNA gene induction is a characteristic of DNA virus infection. Elevated pre-tRNA expression corresponds to increased RNAPIII occupancy for the subset of tRNA genes tested; additionally, post-transcriptional mechanisms contribute to the accumulation of pre-tRNA species. We find increased abundance of tRNA fragments derived from pre-tRNAs upregulated by viral infection, suggesting that non-canonical tRNA cleavage is also affected. Further, pre-tRNA accumulation, but not RNAPIII recruitment, requires gammaherpesvirus-induced degradation of host mRNAs by the virally encoded mRNA endonuclease muSOX. We hypothesize that depletion of pre-tRNA maturation or turnover machinery contributes to robust accumulation of full-length pre-tRNAs in infected cells. Collectively, these findings reveal pervasive changes to tRNA expression during DNA virus infection and highlight the potential of using viruses to explore tRNA biology.

**Significance:** Viral infection can dramatically change the gene expression landscape of the host cell, yet little is known regarding changes in non-coding gene transcription by RNA polymerase III (RNAPIII). Among these are transfer RNAs (tRNAs), which are fundamental in protein translation, yet whose gene regulatory features remain largely undefined in mammalian cells. Here, we perform the first genome-wide analysis of tRNA expression changes during viral infection. We show that premature tRNAs accumulate during infection with the model gammaherpesvirus MHV68 as a consequence of increased transcription, but that transcripts do not undergo canonical maturation into mature tRNAs. These findings underscore how tRNA expression is a highly-regulated process and that cells have strategies to balance tRNA pools during conditions of elevated RNAPIII activity.

## Introduction

The elegant design of the tRNA decoder, an adaptor molecule which converts genetic information into protein, was proposed over 60 years ago (1). Although tRNAs were the first non-coding RNA described, our understanding of tRNA biology, especially their functions outside of protein translation and what dictates their expression, remains limited. This is partly due to the difficulties in applying RNA sequencing and analysis platforms to tRNAs, but recent methodological advances have resulted in a surge of new studies that help to resolve these complexities (2-4). Non-canonical functions of tRNA transcripts have been described, namely the discovery that both premature and mature tRNAs undergo fragmentation into shorter RNAs with numerous regulatory functions in response to stress, cancer, and viral infection (reviewed in (5, 6)). Given the evidence that the ∼400 predicted tRNA genes in mammalian genomes are differentially expressed across cell lines (7-9) and that viral infection can lead to post-transcriptional modulation of tRNAs (10, 11), defining the tRNAome under varying conditions will be central to advancing our understanding of tRNA function, canonical and otherwise.

RNA polymerase III (RNAPIII) transcribes tRNA genes and exhibits enhanced activity during infection with DNA viruses (12-17), many of which encode their own RNAPIII genes. Though RNAPIII activation stimulates expression of viral RNAPIII genes, concomitant accumulation of host RNAPIII transcripts can trigger antiviral immune responses. For example, the induction or misprocessing of RNAPIII-generated host 5S RNA pseudogene and vault RNA transcripts during Herpes Simplex Virus-1 (HSV-1) and Kaposi’s sarcoma-associated herpesvirus (KSHV) infection, respectively, activates RNA sensing pattern recognition receptors (18, 19). The RNAPIII-driven B2 SINE family of retrotransposons are also induced and sensed during infection with the murine gammaherpesvirus MHV68, although MHV68 has uniquely evolved to hijack components of the antiviral NF-κB signaling pathway to instead promote viral gene expression and replication (20-24). While it is clear that induced RNAPIII transcripts can have functional consequences during infection, little is known about the mechanisms underlying the selectivity of RNAPIII activation, including identification of the specific RNAPIII genes induced by viral infection on a genome-wide scale.

DNA viruses upregulate RNAPIII transcription by increasing the abundance of limiting RNAPIII transcription factor complexes specific for certain RNAPIII promoter types (reviewed in (25)). RNAPIII genes have one of three different promoters that recruit a unique combination of the transcription factor complexes, TFIIIA (Gtf3a), TFIIIB (TBP, Brf1 or Brf2, and Bdp1), TFIIIC (Gtf3c1-6) or SNAPc (Snapc1-5) plus RNAPIII for gene expression (26). tRNAs use the abundant RNAPIII promoter type II, which consists of internal A and B boxes that bind TFIIIC and ultimately recruit TFIIIB and RNAPIII to initiate transcription. 7SL, short interspersed nuclear elements (SINEs), vault RNAs, and viral RNAPIII genes also use type II promoters. 5S rRNA genes have an internal type I promoter, which binds TFIIIA (Gtf3a) and requires TFIIIC, TFIIIB, and RNAPIII for transcription. In contrast, type III promoters found in U6 and 7SK genes have external promoters that are TFIIIC-independent and bind the SNAPc complex and a unique TFIIIB complex containing Brf2 protein, rather than Brf1. Defining virally-induced RNAPIII transcript classes and/or transcription factor complexes has been essential for understanding how viruses enhance RNAPIII activity.

The breadth of tRNA function in cells and the fact that many DNA viruses increase RNAPIII activity suggest that regulation of tRNA expression may be a significant point of control during infection. The repertoire of differentially expressed tRNA genes during DNA virus infection is not known, although individual tRNA transcripts are upregulated during EBV and SV40 infection (12, 27). Here, we present the first genome-wide analysis of how infection alters tRNA abundance using the model gammaherpesvirus MHV68. Using a tRNA-specific sequencing method (DM-tRNA-seq) and analysis pipeline, we found that lytic MHV68 infection leads to a ≥3-fold increase in pre-tRNA levels for approximately 14% of tRNA genes. Notably, infection-induced pre-tRNAs do not appear to undergo canonical maturation, as mature tRNA levels remain largely constant, although upregulated pre-tRNAs can be further cleaved into shorter tRNA fragments (tRFs). The accumulation of pre-tRNAs occurs through a combination of transcriptional induction, as evidenced by increased RNAPIII occupancy at activated loci, and post-transcriptional mechanisms. In particular, pre-tRNA accumulation is linked to the activity of the viral endonuclease muSOX/ORF37, which cleaves host mRNAs and downregulates protein synthesis during infection in a process called host shutoff. Given that muSOX is not required for RNAPIII recruitment to tRNA genes, we instead hypothesize that degradation of host mRNAs depletes factors involved in tRNA processing and turnover, contributing to the accumulation of pre-tRNAs. The alterations in tRNA expression during MHV68 infection described here highlight the potential of using viruses to explore mechanistic details of tRNA maturation and turnover.

## Results

### Pre-tRNAs increase in abundance in MHV68-infected cells

Given that MHV68 infection upregulates RNAPIII-driven B2 SINEs (20), which are evolutionarily related to tRNAs, we hypothesized that gammaherpesviral infection may also alter cellular tRNA expression. We first measured pre-tRNA levels by RT-qPCR, using primer sets that were specific for premature tRNAs (pre-tRNAs) produced from intron-containing tRNA-Tyr and tRNA-Leu genes (Figure 1A). During MHV68 infection of NIH3T3 mouse fibroblasts, pre-tRNA levels increased by greater than 15-fold (Figure 1A). This induction appeared specific to tRNAs, as other RNAPIII transcripts (*7SK, U6, 5S*, and *7SL*) did not show differential expression during infection. This is in contrast to Epstein-Barr virus, a gammaherpesvirus that is associated with increased expression of many RNAPIII gene classes due to upregulation of TFIIIB/TFIIIC transcription factor complex subunits at both the mRNA and protein level (12). Accordingly, we did not detect changes in TFIIIB/C subunit mRNA levels upon MHV68 infection (Supplemental Figure 1), and in fact these transcripts were less abundant during infection. This is in line with the general downregulation of host mRNA transcripts that occurs during MHV68-induced host shutoff (28). A time course of MHV68 infection showed that pre-tRNA-Tyr and pre-tRNA-Leu levels were increased by 12 hours post infection (hpi), with their levels accumulating through late stages of infection at 24 hpi (Figure 1B). Notably, the timing of pre-tRNA induction corresponded well with the induction pattern for B2 SINEs. Finally, we also confirmed that pre-tRNA-Tyr and pre-tRNA-Leu are upregulated by northern blot, a method that does not rely on cDNA production (Figure 1C). Mature tRNA-Tyr transcript levels remained constant during infection, suggesting that infection-induced pre-tRNAs may not be processed into mature tRNAs.

**Figure 1.**
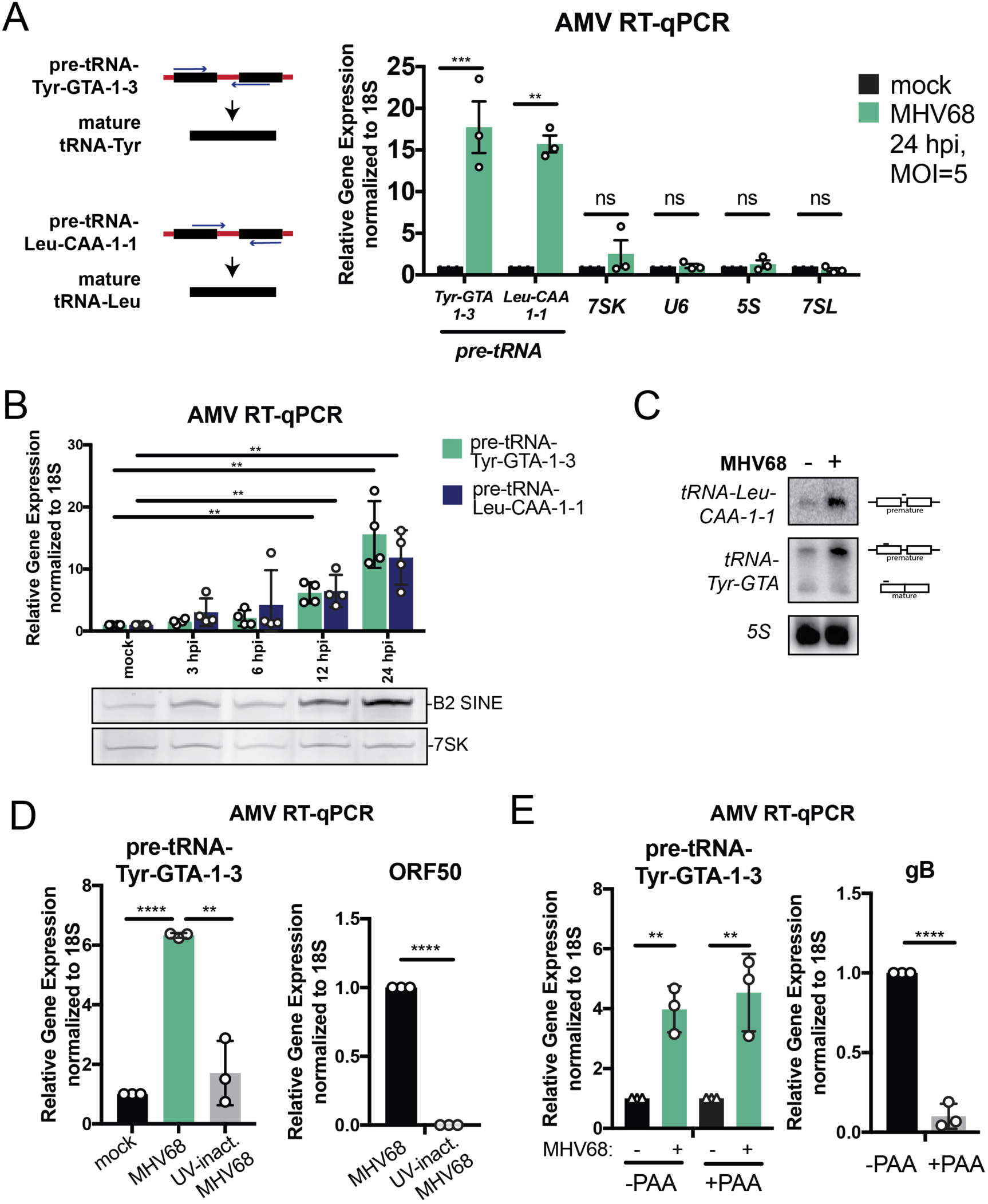
MHV68 infection results in increased pre-tRNA expression. (A) Pre-tRNA-Tyr-GTA-1-3 and pre-tRNA-Tyr-Leu-CAA-1-1 are transcribed with flanking and intron sequences (depicted in red), which are removed during maturation. Pre-tRNAs were detected with forward or reverse primers with 3’ ends complementary to the intron by AMV RT-qPCR. Relative abundance of pre-tRNAs and other RNAPIII transcripts was measured by RT-qPCR in mock and MHV68-infected NIH3T3s. Expression was normalized to 18S rRNA and compared to values from mock-infected cells. (B) RNA was extracted from NIH3T3 cells following a time course of MHV68 infection and used for both AMV RT-qPCR to detect pre-tRNAs and primer extension to detect B2 SINE and 7SK transcripts. (C) Northern blot was performed on RNA from mock or MHV68-infected MC57Gs. Transcripts produced from tRNA-Leu-CAA-1-1 were detected with intron and exon-specific probes. tRNA-Tyr transcripts were detected with a probe that anneals with a conserved 5’ region in the tRNA-Tyr family. Probe binding is depicted on the right. (D) RNA from mock and MHV68-infected MC57Gs +/− UV-inactivation was used for AMV RT-qPCR. ORF50 is an early viral gene. (E) Mock and MHV68-infected MC57Gs were treated with 200 ng/μl phosphonoacetic acid (PAA) to block viral DNA replication, and extracted RNA was used for RT-qPCR analysis. gB is a late viral gene whose expression is dependent on successful viral DNA replication. All infections were done at an MOI=5 for 24 hours unless otherwise noted. RT-qPCR experiments were done in triplicate at a minimum. Error bars show SD and statistics were calculated using an unpaired t-test. *P<0.05, **P<0.01. ***P<0.001.

Given the kinetics of induction, we hypothesized that events occurring mid to late infection were responsible for pre-tRNA accumulation, as opposed to a cellular response to viral binding and entry. In agreement with this hypothesis, infection of MC57G fibroblasts with UV-inactivated MHV68 did not lead to pre-tRNA-Tyr induction (Figure 1D). Additionally, wild type MHV68 infection of the MC57G fibroblast cell line led to an induction similar to NIH3T3s. UV-inactivation prevented post-entry viral gene expression, as demonstrated by the absence of expression of the immediate early viral gene ORF50. To narrow down the gene class responsible for the kinetics of tRNA induction, we performed infections in the presence of phosphonoacetic acid (PAA), which blocks viral DNA replication and prevents the expression of viral late genes, such as gB. Although late gene expression was inhibited, pre-tRNA-Tyr accumulated to similar levels in cells infected in the presence of PAA (Figure 1E). Thus, one or more effects of early viral gene expression underlie pre-tRNA accumulation during MHV68 infection. Together, these data demonstrate that early viral gene expression drives the increased abundance of host pre-tRNAs.

### Dm-tRNA-seq reveals upregulation of premature tRNAs during infection

Extensive base modification of tRNAs can prevent reverse transcription of full-length cDNA products. Although there are some modifications present in pre-tRNAs, they likely have not undergone extensive modification and therefore are more efficiently amplified relative to mature tRNAs during reverse transcription (4, 29). Amplification of full-length cDNA products from tRNAs can be improved by using the highly processive thermostable group II-intron containing reverse transcriptase (TGIRT) (30) and by partial removal of tRNA strong-stop modifications by treating with AlkB demethylases (2, 3). To define the extent of tRNA upregulation genome-wide, we generated sequencing libraries using TGIRT and AlkB-treated small RNA (a method called DM-tRNA-seq (3)) in four biological replicates of mock versus MHV68 infected MC57G cells. Our analysis pipeline was designed to distinguish reads from pre-tRNAs and mature tRNAs, as our preliminary data with tRNA-Leu and tRNA-Tyr suggested that MHV68 may elevate pre-tRNA but not mature tRNA levels (Figure 1C). Reads were first aligned to mature tRNA sequences, which contained predicted tRNA genes from tRNAscan-SE (31, 32) modified by intron removal and the addition of mature –CCA 3’ ends, yielding counts for mature tRNAs. The remaining reads were aligned to a masked genome appended with pre-tRNA sequences including intact introns and 50 nt of upstream and downstream flanking sequence. Thus, classification as a pre-tRNA required the absence of a 3 –CCA and the presence of 5’ leader, intron, or 3’ trailer sequence.

tRNAscan-SE yields a list of genes containing both true tRNAs and tRNA-like sequences such as pseudo-tRNAs and B2 SINE elements. Among these, we focused on the high confidence set of tRNA genes as defined by the genomic tRNA database (n=405) (31, 32). Due to unique genomic sequences found in 5’ leader and 3’ trailers, pre-tRNA reads were less likely to multimap to the genome and were used as a proxy for expression of the respective tRNA genes. Of the 405 predicted high confidence tRNA genes, we were able to map pre-tRNAs originating from 313 tRNA genes using Salmon (Supplemental File 1) (33). Additionally, we used a different mapping strategy analyzing only the uniquely mapping reads in order to more confidently identify the locus of origin for expressed pre-tRNAs, although this strategy may underestimate the total number of expressed pre-tRNAs. Restricting the data to uniquely mapping reads yielded a total of 244 tRNA genes with detectable pre-tRNAs from infected cells.

Differential gene expression analysis revealed that pre-tRNAs are more abundant during MHV68 infection (Figure 2A, blue and orange circles). In fact, raw read count ratios alone (sum of total reads per replicate compared to mock) displayed an increase in pre-tRNA reads present in infected cells compared to uninfected cells (Figure 2B). It is likely that many of the highly expressed pre-tRNA genes (Figure 2A, blue circles) that are not in the high confidence tRNA gene list (Figure 2A, orange circles) are SINE retrotransposons, as B2 SINEs are robustly induced during infection (20, 34). Focusing on high confidence tRNA genes, we found many of these loci produce more pre-tRNAs during infection (Figure 2A, orange circles; Figure 2C). Yet, while total levels of pre-tRNAs were higher in infected cells, we observed selectivity in which tRNA genes were upregulated. Of the high confidence pre-tRNAs detectably expressed in uninfected MC57G cells, approximately one-fifth were significantly upregulated during MHV68 infection, depending on the mapping strategy used (Figure 2C shows results with Salmon: 43 out of 313 (13.7%), unique mappers: 63 out of 244 (25.8%), FDR<0.05). However, a general trend of increased expression of pre-tRNAs was observed as judged by a fold change greater than 2 (Figure 2C shows results with Salmon: 141 out of 314 (44.9%); unique mappers: 158 out of 244 (64.8%)). In general, at least one member of each tRNA isotype (tRNAs encoding the same amino acid) were present in the differential hits, but not all isodecoders (tRNAs with the same anticodon) were represented (Figure 2D, Supplemental Figure 2). A triple A box motif that is present in each enriched in the differential hits (Supplemental Figure 3).

**Figure 2.**
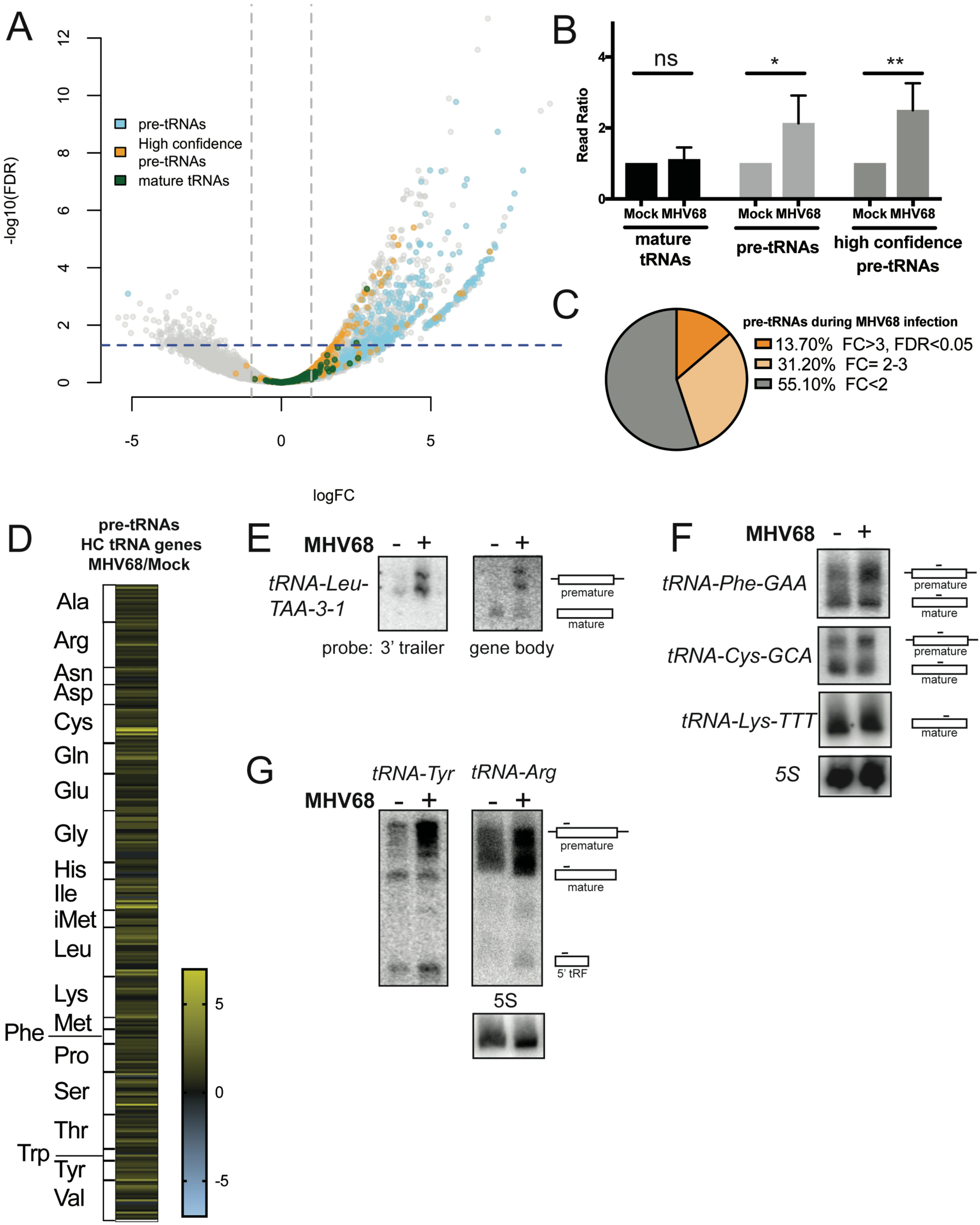
DM-tRNA-seq reveals differential expression of the pretRNAome upon infection. (**A**) Differential gene expression analysis from DM-tRNA-seq was plotted as fold change (FC) versus the false discovery rate (FDR). Pre-tRNAs (high confidence in yellow, other in blue), mature tRNAs (high confidence only in red), and other small RNA reads (gray) are annotated and the dotted lines represent the boundaries of FC=2 and FDR=0.05. High confidence refers to the tRNA gene set predicted by GtRNAdb. (**B**) Raw read counts for mature tRNAs, pre-tRNAs, or high confidence pre-tRNAs from DM-tRNA-seq of MHV68-infected MC57Gs were compared to those of uninfected cells. Error bars show SD and statistics were calculated using an unpaired t- test on raw read counts from four replicates. (**C**) A pie chart shows the percentage of high confidence pre-tRNAs detected by DM-tRNA-seq that were significantly upregulated (dark orange, FC>3 and FDR<0.05) versus upregulated but not significant (light orange, FC>2 and FDR>0.05) versus unregulated (gray, FC<2). (**D**) A heat map of log_2_FC changes of pre-tRNA levels by DM-tRNA-seq by isotype. (**E**) RNA from mock or MHV68-infected MC57Gs was used for northern blot using probes specific for either the 3’ trailer or gene body of tRNA-Leu-TAA-3-1. (**F**) RNA from mock or MHV68-infected MC57Gs was used for northern blot using probes specific for the gene body of the tRNA-Phe-GAA, tRNA-Cys-GCA, and tRNA-Lys-TTT families. Probes detected tRNAs from multiple loci. (**G**) RNA from mock or MHV68-infected MC57Gs was used for northern blot using probes specific for the 5’ exon of tRNA-Tyr-GTA and tRNA-Arg-TCT families. Probes detected tRNAs from multiple loci. All infections were done in MC57G fibroblasts at an MOI=5 for 24 hours.

Unlike pre-tRNAs, the levels of most mature tRNAs did not change in a statistically significant manner, suggesting that the upregulated pre-tRNAs do not undergo canonical maturation (Supplemental File 1). In fact, only two high confidence tRNA genes (tRNA-Lys-TTT-2-2 and tRNA-Leu-TAA-3-1) showed significant differential expression at the level of the mature tRNA transcript (Figure 2A, green circles FDR<0.05). To validate these findings, we designed northern blot probes to the 3’ trailer or gene body specific to transcripts originating from tRNA-Leu-TAA-3-1 (Figure 2E). While pre-tRNA transcripts were elevated from the tRNA-Leu-TAA-3-1 gene in infected cells, levels of mature tRNAs did not change upon infection. We were not able to design a probe specific for tRNA-Lys-TTT-2-2 transcripts, but using a probe that recognizes members of the tRNA-Lys-TTT family did not show any differences in mature tRNA-Lys-TTT expression during MHV68 infection (Figure 2F). Similarly, other tRNA families that we tested by northern blotting, including tRNA-Phe-GAA and tRNA-Cys-GCA families, showed upregulation only at the level of the full-length pre-tRNA (Figure 2F).

### MHV68 induces differential expression of pre-tRNA fragments

Studies have demonstrated that differential expression of tRNAs, either when comparing different mammalian cell types, or upon deletion of the RNAPIII negative regulator, Maf1, is not associated with major changes in steady state mature tRNA levels (36, 37). Instead, increased tRNA expression largely seems to affect pre-tRNA and/or tRNA fragment (tRF) levels. DM-tRNA-seq analysis revealed increased expression of pre-tRNAs, but the size selection we used for this pipeline (50-200 nt) did not allow for interrogation of tRF expression. To investigate whether MHV68 infection leads to changes in tRF levels, we performed northern blot analysis using probes that detect Tyr or Arg tRFs derived from the 5’ end of the pre-tRNA transcripts, previously reported in mouse embryonic fibroblasts (38) (Figure 2G). We found that Tyr and Arg tRF abundance was modestly increased upon infection with MHV68.

The reported activities for tRFs are strikingly diverse, rendering functional predictions difficult, particularly under conditions such as infection where a multitude of different fragments may be produced (11, 39). Nonetheless, we explored possible activity for the Tyr tRF, as its overexpression has been previously reported to enhance p53 activation through phosphorylation at serine 18 (S18) in response to oxidative stress (38). Because MHV68 infection also leads to p53 activation through phosphorylation at S18 (40), we examined whether this phenotype was linked to pre-tRNA induction. However, depletion of the TFIIIB subunit Brf1 required for Pol III activity did not alter the p53 phosphorylation status in MHV68 infected cells (Supplemental Figure 4). Thus, while MHV68-induced pre-tRNAs may undergo processing into small RNA fragments, whether these harbor specific functions (for example under specific cell contexts) remains unknown. Regardless of their function, virus-induced pre-tRNA accumulation provides a platform for dissecting the mechanisms underlying tRNA gene expression control.

### Increased levels of RNA Polymerase III are found at tRNA genes associated with increased pre-tRNA expression during infection

The distinct subset of tRNA genes that have higher expression at the pre-tRNA level during MHV68 infection could be reflective of increased transcription, stability, or a combination of both. To determine if the increase in pre-tRNA transcripts during infection results from transcriptional upregulation, we performed chromatin immunoprecipitation followed by qPCR (ChIP-qPCR) to measure occupancy of the RNAPIII subunit Polr3A at several loci in mock and MHV68-infected MC57G fibroblasts. RNAPIII abundance specifically increased during MHV68 infection at each of the tested differentially expressed loci (tRNA-Tyr-GTA-1-3, tRNA-Leu-CAA-1-1, and tRNA-Leu-TAA-3-1) but remained unchanged at a tRNA gene not differentially expressed (tRNA-Ile-TAT-2-3) and at 7SK (Figure 3A). RNAPIII abundance at the RNAPII-dependent GAPDH promoter was included as a negative control to show specificity of the RNAPIII ChIP signal. These data suggest that there is an increase in RNAPIII recruitment to select tRNA genes during MHV68 infection.

**Figure 3.**
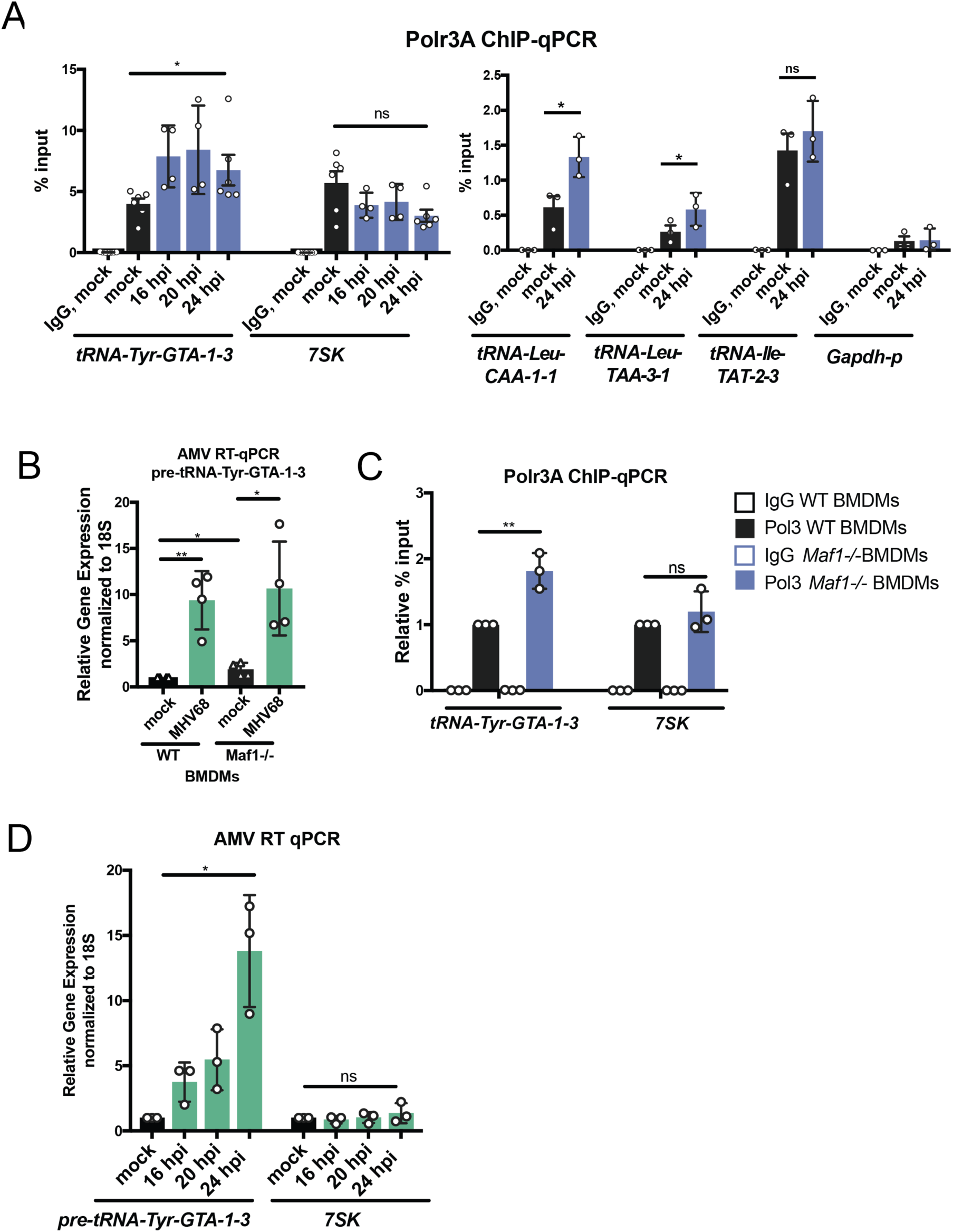
Polr3A is recruited to induced tRNA loci in infected cells. (A) RNAPIII Occupancy was measured in mock and MHV68-infected MC57Gs using ChlP-qPCR with antibodies against the Polr3A subunit at three tRNA genes upregulated at the pre-tRNA level by DM-tRNA-seq (tRNA-Tyr-GTA-1-3, tRNA-Leu-CAA-1-1, tRNA-Leu-TAA-3-1), a tRNA gene not associated with increased pre-tRNA levels (tRNA-Ile-TAT-2-3), 7SK, and the promoter of RNAPII-driven Gapdh. (B) RT-qPCR was performed with RNA extracted from wild type and Maf1-/-MHV68-infected bone marrow derived macrophages (BMDMs). (C) RNAPIII Occupancy was measured using Polr3A ChlP-qPCR on wild type and Maf1 -/- BMDMs at tRNA-GTA-1-3, a tRNA gene also upregulated during MHV68 infection, and 7SK. Results are reported in relative % input. (D) RNA from mock or MHV68-infected MC57Gs was extracted at the indicated hours post infection (hpi) and subjected to RT-qPCR. All infections were done in MC57G fibroblasts at an MOI=5 for 24 hours except where indicated. Error bars show SD and statistics were calculated using an unpaired t-test (RT-qPCR) or paired t-test on raw % input values (ChlP-qPCR). *P<0.05, **P<0.01. ***P<0.001.

Maf1 is a well-studied negative regulator of transcription by RNAPIII whose activity is controlled by phosphorylation and cellular localization in response to nutrients and stress conditions (41, 42). When active, Maf1 directly targets the Brf1 subunit of TFIIIB and RNAPIII to inhibit transcription. In addition, in proliferating and differentiated mammalian cells, Maf1 chronically represses RNAPIII transcription to balance RNA synthesis with cellular demand (43, 44). Thus, if the mechanism underlying MHV68 induction of pre-tRNAs involved antagonizing Maf1-dependent repression, we reasoned that cells lacking Maf1 should no longer display a difference in pre-tRNA levels between uninfected and infected cells. To test this hypothesis, we derived primary bone marrow macrophages (BMDM) from WT and *Maf1-/-* mice and infected them with MHV68 at an MOI=5. MHV68 infection of BMDMs resulted in ∼10-fold increase in pre-tRNA-Tyr upon MHV68 infection, demonstrating that increased pre-tRNA expression during infection occurs in primary cells and is not unique to fibroblast cell lines (Figure 3B). MHV68-infected WT and *Maf1-/-* BMDMs showed similar levels of pre-tRNA expression, indicating that viral induction of pre-tRNA-Tyr levels was not dependent on *Maf1* regulation of RNAPIII. Similar to other published reports (36), uninfected *Maf1*-/-cells showed a ∼2-fold increase in pre-tRNA-Tyr compared to WT BMDMs as well as increased RNAPIII promoter occupancy in ChIP assays, confirming that RNAPIII activity is elevated in the absence of this negative regulator (Figure 3B, 3C).

Even though the relative increases in pre-tRNA-Tyr levels were dramatically larger in MHV68-infected cells compared to cells lacking Maf1 (Figure 3B,D), we were struck by the observation that the increase in RNAPIII occupancy in these two scenarios was comparable (Figure 3A,C). To explore the contribution of RNAPIII recruitment to pre-tRNA levels, we compared pre-tRNA abundance by RT-qPCR (Figure 3D) and RNAPIII promoter occupancy by Polr3A ChIP-qPCR (Figure 3A) across a time course of MHV68 infection in MC57G fibroblasts. Pre-tRNA-Tyr levels were induced about 5-fold at 16 and 20 hpi but increased to a nearly 15-fold induction at 24 hpi, whereas *7SK* levels remained constant (Figure 3D). In contrast to the RNA abundance data, RNAPIII occupancy at the pre-tRNA-Tyr promoter was modestly increased by 16hpi, but did not further increase at later time points (Figure 3A). Thus, after 16 hpi, it is likely that post-transcriptional mechanisms underlie the continued accumulation of pre-tRNA-Tyr.

### Virus-induced mRNA degradation contributes to post-transcriptional accumulation of pre-tRNAs

We reasoned that selective post-transcriptional accumulation of pre-tRNAs might occur if pre-tRNA maturation or turnover were impaired, for example if the virus down-regulated factors involved in these processes. A well-characterized phenotype of MHV68 infection is the widespread degradation of host mRNAs, which is caused by the viral mRNA-specific endonuclease muSOX (28). Notably, the levels of the RNAPIII-transcribed viral tRNAs (vtRNAs) are decreased in cells infected with the MHV68 muSOX point mutant R443I (MHV68.R443I), which has reduced host shutoff activity (45). muSOX is expressed with early kinetics, but its mRNA depletion phenotype is most prominent at late stages of infection, coincident with maximal pre-tRNA accumulation. We tested whether virus-induced mRNA decay contributes to post-transcriptional accumulation of cellular pre-tRNAs by measuring pre-tRNA abundance in MC57G cells infected with MHV68.R443I or the mutant revertant (MR) virus in which the R443I was restored to wild type (Figure 4A). Similar to the vtRNAs, the levels of cellular pre-tRNA-Tyr and pre-tRNA-Leu were lower levels in cells infected with the R443I mutant compared to MR MHV68 (Figure 4A). In contrast, the viral ORF50 mRNA was slightly increased during the R443I mutant infection, consistent with previous reports (45). We verified these results using northern blot analysis (Figure 4B). To determine if these changes reflected transcriptional output, we measured RNAPIII binding at two differentially regulated host tRNA genes and the viral TMER2 locus during MR vs R443I infection (Figure 4C). RNAPIII abundance was similar during infection with MR and R443I viruses, in stark contrast to the difference in pre-tRNA and vtRNA transcript abundance. This result suggests that the decreased abundance of these transcripts in total RNA is not due to decreased levels of transcription. Collectively, these data reveal that the mRNA decay phenotype associated with WT MHV68 infection is not involved in transcriptional activation of tRNA loci, but instead stabilizes the unprocessed pre-tRNAs.

**Figure 4.**
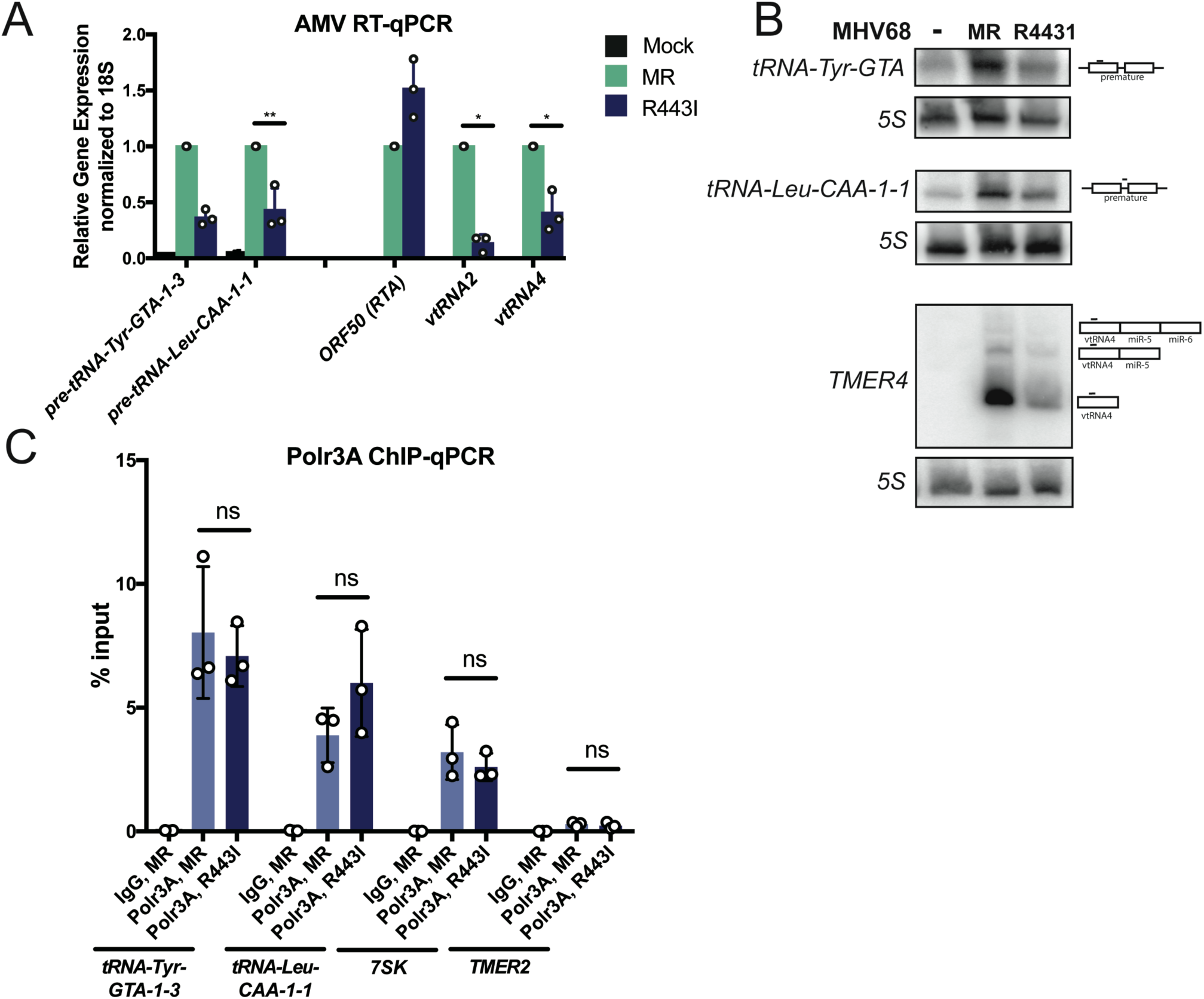
muSOX activity is required for pre-tRNA accumulation, but not for RNAPIII recruitment. **(A)** MC57Gs were infected with muSOX R443I or mutant revertant (MR) MHV68 and RT-qPCR was performed with extracted RNA. (B) Northern blots were performed on RNA from the same conditions as in (A) using probes to detect pre-tRNA-Tyr, pre-tRNA-Leu-CAA-1-1, and viral TMER4 transcripts. The TMER4 probes bind the vtRNA4 sequence. (C) RNAPIII occupancy was measured using ChIP-qPCR with antibodies against the PolrSAsubunit of RNAPIII at induced tRNA loci, viral TMER2, and 7SK using the cells described in (A). All infections were done at an MOI=5 for 24 hours. Error bars show SD and statistics were calculated using an unpaired t-test on raw ΔCt values (RT-qPCR) or paired t-test on raw % input values (ChIP-qPCR). *P<0.05, **P<0.01. ***P<0.001.

## Discussion

This study represents the first genome-wide analysis of how infection modulates the tRNAome, revealing that the model gammaherpesvirus MHV68 causes extensive changes to pre-tRNA but not mature tRNA abundance. Additionally, we found evidence that elevated levels of pre-tRNA-Tyr and -Arg leads to an increase in production of their cognate tRFs. We did not detect increased expression across all tRNA genes, but instead found a dramatic accumulation of pre-tRNAs from a subset of loci, which may reflect differences in promoter accessibility, regulation and/or pre-tRNA processing. Collectively, our data support a model whereby pre-tRNA accumulation during MHV68 infection is driven by both RNAPIII recruitment and RNA stabilization (Figure 5). We hypothesize that early events in infection stimulate RNAPIII-driven transcription of specific tRNA loci, while muSOX-induced depletion or relocalization of pre-tRNA processing machinery stalls further canonical processing of these transcripts, leading to their build up late in infection (Figure 5). These findings are bolstered by early reports showing enhanced RNAPIII activity during DNA virus infection, including for select tRNAs (12-17, 27), suggesting that manipulation of the pre-tRNA landscape is likely to be a conserved feature of numerous viruses.

**Figure 5.**
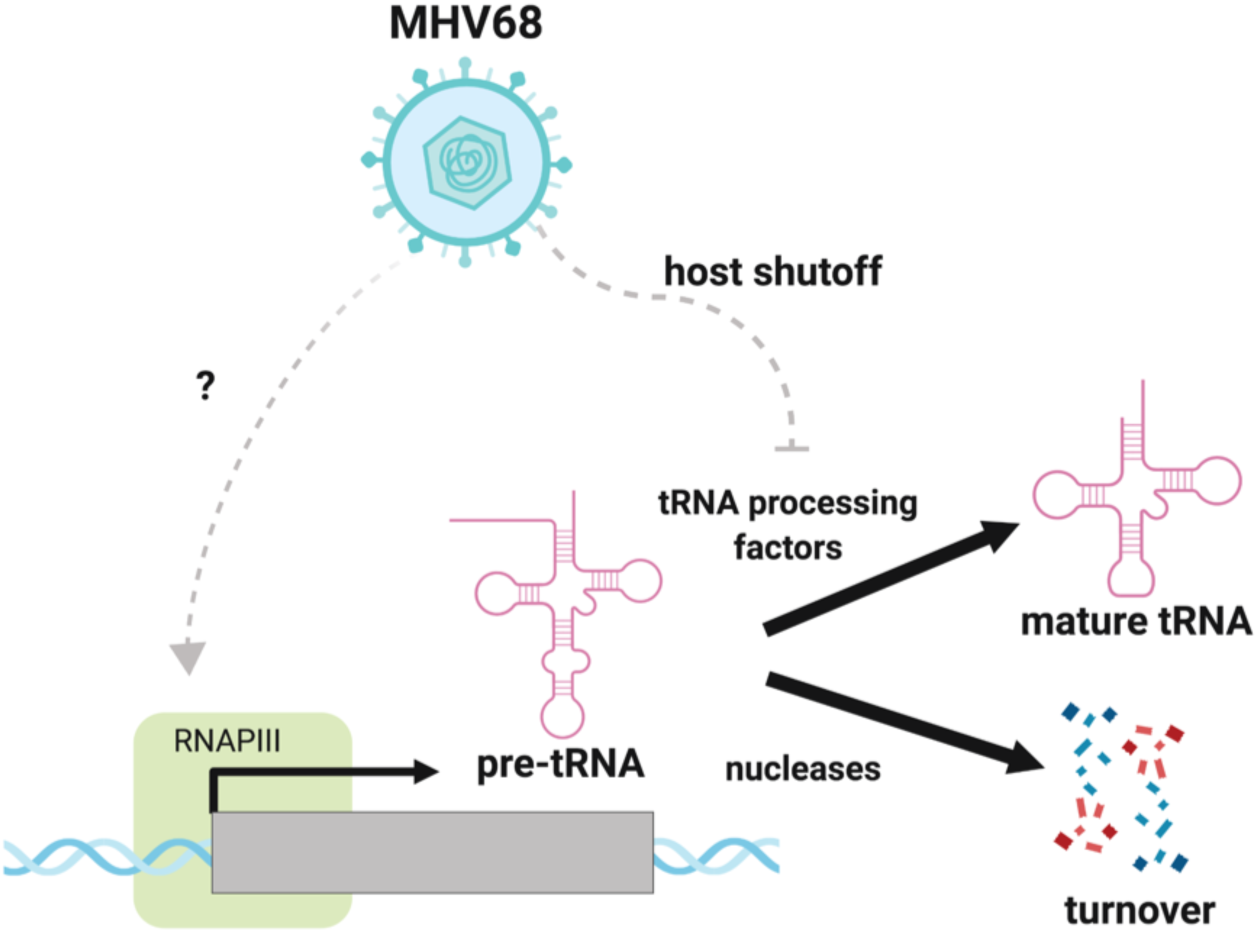
MHV68 infection induces pre-tRNA accumulation by transcriptional and post-transcriptional mechanisms. RNAPIII is recruited to specific tRNA genes by a Maf1-Independent mechanism, increasing the output of pre-tRNAs. Pre-tRNAs accumulate in infected cells due to depletion of maturation or turnover machinery during muSOX-mediated mRNA decay (host shutoff).

Pre-tRNA steady state levels are dependent on both the production and stability of the pre-tRNA transcripts. We performed Polr3A ChIP-qPCR and found that increased pre-tRNA abundance was associated with higher levels of Polr3A occupancy during infection. In contrast, a tRNA gene with constant pre-tRNA expression did not show changes in Polr3A binding. This confirms that transcriptional increases – at least at some tRNA genes – contribute to pre-tRNA accumulation during infection. Whether this is true for all tRNA genes associated with increased pre-tRNA levels remains to be determined. Additionally, increased RNAPIII occupancy during infection suggests a mechanism involving RNAPIII recruitment to specific tRNA genes, and raises the possibility that these tRNA genes are not expressed to their full potential in uninfected cell populations. In support of this, there is considerable overlap between tRNA genes upregulated during MHV68 infection with those upregulated in cells lacking Maf1, a negative regulator of RNAPIII that prevents polymerase recruitment (59 of 79 pre-tRNAs induced by MHV68 infection of MC57G fibroblasts were also increased in *Maf1-/-* versus WT liver in (36)). However, Maf1 does not appear to play a pivotal role in pre-tRNA accumulation during MHV68 infection, as the increase in pre-tRNA abundance does not require Maf1.

The mechanisms employed during MHV68 infection to increase RNAPIII recruitment have not been identified. During EBV infection, TFIIIC subunit and Bdp1 protein levels increase, along with transcripts for RNAPIII genes with type I and II promoters (*7SL, 5S*, tRNAs), but not those with type III promoters (*7SK, U6, MRP*) (12). In contrast, MHV68 infection leads to transcriptional induction of B1/B2 SINEs (20, 34) and certain tRNA genes, all of which have type II promoters. TFIIIB/C subunit mRNA levels do not increase during MHV68 infection, and quantitative mass spectrometry of MHV68-infected cells indicates that total TFIIIB/C protein levels are not elevated, although there may be enrichment of these factors in the nucleus (46). Thus, RNAPIII regulation during MHV68 infection appears distinct from that observed with EBV and may therefore involve novel mechanisms. Like EBV, SV40 infection leads to increased TFIIIC protein levels, but additionally results in higher availability of TFIIIB by large T-antigen sequestration of the retinoblastoma tumor suppressor protein (RB), a negative regulator of RNAPIII transcription (47). The changes in RNAPIII output in mouse fibroblasts are similar between SV40 and MHV68 infections (B1/B2 SINEs and certain tRNAs are induced (27, 48)); additionally, mass spectrometry data suggests that RB subcellular concentration changes during MHV68 infection (46). Thus, future studies will explore how RB is regulated during MHV68 infection and whether this protein is an important player in gammaherpesvirus-induced RNAPIII activation. Potential roles of RB could include the posttranslational regulation of transcription factor activities, altered factor availability or epigenetic changes influencing gene accessibility.

With the exception of selenocysteine tRNA (7), all tRNAs genes use the same internal A/B box promoter architecture for transcription. However, based on differential binding of RNAPIII, many studies have revealed that tRNA genes are not equally transcribed in mammalian cells, as measured by the differential binding of RNAPIII (7-9). Genome-wide studies examining chromatin accessibility, modification and transcription factor binding have suggested that a driver of RNAPIII occupancy at tRNA genes is open and active chromatin, typically in the vicinity of RNAPII transcription sites, yet the basis for differential RNAPIII binding remains to be determined (7, 8, 49, 50). Further studies are needed to assess how host chromatin accessibility and modifications change during MHV68 infection. Interestingly, MHV68 infection leads to depletion of RNAPII-specific subunits, resulting in less RNAPII transcription of the host genome during infection (46). There is some evidence of transcriptional interference between RNAPIII and RNAPII bound at nearby genes, as depletion of the Rbp1 subunit of RNAPII leads to upregulation of a subset of tRNA genes (51). It is possible that depletion of RNAPII allows for increased recruitment of RNAPIII machinery. We hypothesize that a combination of genome accessibility and transcription factor availability are associated with tRNA transcriptional increases during infection.

Although tRNA gene transcription is elevated due to RNAPIII recruitment, the magnitude of pre-tRNA abundance is unlikely to be explained by transcriptional activation alone. Instead, our data suggest that pre-tRNA stability is also increased. While we have not been able to measure pre-tRNA half-life directly (due to difficulty in robustly inhibiting RNAPIII in the required short time frame of pre-tRNA processing), it is clear that pre-tRNAs accumulate dramatically between 20-24 hpi, a time frame in which RNAPIII occupancy is unchanged. Infection-induced pre-tRNAs do not seem to be processed into mature tRNAs, as we found no evidence of differential expression at the mature tRNA transcript level. Similarly, *Maf1-/-* tissues exhibit upregulation of pre-tRNAs without significant changes to mature tRNA levels, suggestive of a homeostatic mechanism to maintain the size of mature tRNA pools when tRNA gene transcription is increased (36). In both of these cases, it is likely that the increase of pre-tRNAs upon transcriptional induction saturates processing machinery, an outcome well established in yeast (52, 53). It was striking that some pre-tRNA transcripts revealed in our study were much more highly expressed than others. These pre-tRNAs might have specific modifications, unique structural characteristics, or processing fates that render them inherently more stable in the cell or during infection. Additionally, excess pre-tRNAs might be cleaved into stable, and perhaps functionally distinct, fragments. We demonstrated here one example of a tRF derived from pre-tRNA-Tyr that is elevated during MHV68 infection. This Tyr tRF was originally described to be abundant in the absence of the Clp1 kinase, a known regulator of the TSEN tRNA splicing complex (38, 54, 55). As the 3’ end of this tRF corresponds to the exact end of the 5’ exon boundary (38), it is possible that the Tyr and Arg tRFs are splicing intermediates and thus constitute evidence that splicing of pre-tRNA transcripts is indeed perturbed upon infection. However, as only a small fraction of pre-tRNAs undergo splicing, there are likely many aspects of pre-tRNA maturation that are perturbed in the infection context. Regardless, knowing the identity of hyperabundant pre-tRNAs will help guide studies to investigate unique features and broaden our understanding of tRNA maturation and turnover.

We hypothesized that tRNA processing and turnover machinery might be depleted in infected cells due to gammaherpesvirus-induced host shutoff implemented at late stages of viral infection. Widespread mRNA decay is initiated by a gammaherpesvirus-encoded endonuclease, called SOX in KSHV and muSOX in MHV68. SOX and its homologs cleave RNAPII-transcribed mRNAs, leading to subsequent downregulation of host protein synthesis (28, 56). Depletion of host proteins might further increase the ratio of pre-tRNA to available processing and turnover machinery, leading to increased pre-tRNA half-lives. Though total protein levels of tRNA processing machinery are not dramatically altered during a 24-hour lytic infection cycle, there are some decreases in nuclear abundance of tRNA-related factors during infection, including the nuclear exosome (46). Modest abundance changes of many tRNA-related processing and turnover complexes could have an additive effect on the stability of pre-tRNAs in infected cells. Moreover, pre-tRNAs were less abundant in cells infected with the host shut-off defective R443I virus, a phenotype not explained by changes in Polr3A abundance at tRNA genes. Viral TMER transcripts are also less abundant under these conditions (45), suggesting that the factors involved with pre-tRNA stability during infection may also be involved in viral TMER transcript stability. TMERs consist of a tRNA-like sequence with 1-2 miRNA stem loops that are processed by the host tRNA 3’ processing enzyme tRNAse Z (57, 58). Whether tRNAse Z or other host factors are involved in stabilizing host pre-tRNA and viral TMERs is an important question for future studies.

Boosting RNAPIII activity is thought to promote the expression of RNAPIII-transcribed genes encoded by some DNA viruses (e.g., Epstein-Barr virus EBERs, Adenoviruses VA RNAs, MHV68 TMERs) -- however, not all DNA viruses encode RNAPIII genes, suggesting that regulation of RNAPIII could be advantageous to the virus or host for other reasons. For example, because nascent RNAPIII transcripts contain 5’ triphosphate ends, they can and do serve as substrates for RNA sensors in the cell, such as RIG-I (18, 19). These studies show that while typically the 5’ ends of RNAPIII transcripts are typically processed or protected from sensing by protein binding partners, HSV-1 and KSHV infection tips the balance and results in exposure of 5’ triphosphate ends of RNAPIII transcripts for RIG-I recognition. Additionally, increased tRNA fragmentation observed during MHV68 infection could result in modulation of gene regulation, cell proliferation, and stress responses, as has been reported in for tRFs produced during viral infection or cancer (5, 6). A thorough examination of tRFs produced during gammaherpesvirus infection by small RNAseq will be required to begin to address the role of tRF generation upon infection. In summary, exploring viral modulation of host tRNAomes could lead to exciting insights both for tRNA biology and for the role of RNAPIII activation during DNA virus infection.

## Materials and Methods

### Cell and Virus Culture

NIH3T3 and MC57G cell lines were obtained from ATCC, maintained in Dulbecco’s modified Eagle medium (DMEM; Gibco) supplemented with 10% fetal bovine serum (FBS; VWR), and screened regularly for mycoplasma by PCR. Wild-type and *Maf1* knockout primary bone marrow-derived macrophages (BMDMs) were differentiated from adult 3-6 month old C57BL/6 mice (36). Provided femurs and tibia were ground in a sterile tissue-culture hood using a mortar and pestle in DMEM before being passed through a 70 μm filter to remove debris. Cells were differentiated in macrophage media (High glucose DMEM, 10% FBS, 10% MCSF, 1% GlutaMax) + 1% PenStrep for 7 days before being frozen for later use. After thaw, cells were not subjected to any media containing antibiotics. MHV68 was amplified in NIH3T12 cells and TCID50 was measured on NIH3T3s by limiting dilution. GFP-expressing wild type (59) and GFP-expressing ΔHS (SOX R443I (60)) MHV68 viruses were incubated with cells for 1 hour at an MOI of 5 to allow viral entry, and then the virus-containing media was removed and replaced with fresh media. PAA was used at a concentration of 200 μg/ml and was added at the start of the infection. UV-inactivation of MHV68 was performed by autocrosslinking twice in a Stratalinker 2400 (1200 μJ X 100). For infection of BMDMs, virus was added to cells in serum-free DMEM for 4 hours in non-TC treated plates. Virus containing media was then aspirated and replaced with macrophage media without antibiotics.

### RNA isolation and analysis

Total RNA was isolated from cells using TRIzol (Invitrogen), treated with Turbo DNase (Ambion) and reverse transcribed with AMV RT (Promega) primed with random 9-mers. For TGIRT-qPCR, cDNA was synthesized with the TGIRT™-III Enzyme (InGex) following manufacturer’s instructions. Quantitative PCR analysis was performed with iTaq Universal SYBR Green Supermix (Bio-Rad) using the primers listed in Supplemental Table 1. qPCR was performed on at least three biological replicates and Ct values were measured from three technical replicates per biological sample. Fold change was calculated by ΔΔCT method. For northern blots, 20 μg of RNA was loaded into 8-12% PAGE/7 M Urea gels, transferred to Hybond-XL or N+ (GE) membranes using a Trans-blot Turbo Transfer system (Bio-Rad). Blots were pre-hybridized in ULTRAhyb buffer (Thermo Fisher) at 42 °C for 1 hour before adding radiolabeled probe. Probes were generated by end-labeling oligos listed in Supplemental Table 1 using T4 PNK and [γ-^32^P]-ATP. Blots were probed overnight at 42 °C, washed 3 times for 5 minutes in 0.5x SSC and 0.1% SDS. Primer extension was performed on 10 μg of RNA using a 5’ fluorescein labeled oligo specific for B2 SINEs or 7SK. RNA was ethanol precipitated and resuspended in 9 μl of annealing buffer (10 mM Tris-HCl, pH7.5, 0.3 M KCl, 1 mM EDTA) plus 1 μl of 5’ fluorescein labeled oligo. Sample was briefly heated to 95 °C, then incubated at 55 °C for 1 hour. 40 μl of extension buffer (10 mM Tris-HCl, pH 8.8, 5 mM MgCl_2_, 5 mM DTT, 1 mM dNTP) and 1 μl of AMV RT (Promega) was added and the sample was incubated at 42 °C for 1 hour. Nucleic acids were precipitated, resuspended in RNA loading dye, and separated in 8% PAGE/ 7 M Urea gels.

### DM-tRNA-seq library preparation

Protocol was modified from (3). His-tagged wild-type and D135S AlkB plasmids were obtained from Addgene (#79050 and #79051) and proteins were purified by a Ni-NTA column followed by cation exchange. Total RNA extracted from four biological replicates was spiked with *in vitro* transcribed *E. coli* tRNA-Lys, *E. coli* tRNA-Tyr, and *S. cerevisiae* tRNA-Phe transcripts at 0.01 pmol IVT tRNAs per μg total RNA. RNA was deacylated in 0.1M Tris-HCl, pH 9 at 37 °C for 45 min, ethanol purified, and then dephosphorylated with PNK. Deacylated and dephosphorylated RNAs were then purified with a mirVANA small RNA purification kit (Ambion) to isolate RNAs ≤ 200 nucleotides. Small RNAs were demethylated in 300 mM NaCl, 50 mM MES pH 5, 2 mM MgCl_2_, 50 μM ferrous ammonium sulfate, 300 μM 2-ketoglutarate, 2 mM L-ascorbic acid, 50 μg/ml BSA, 1U/μl SUPERasin, 2X molar ratio of wt AlkB, and 4X molar ratio of D135S AlkB for 2 hours at room temperature. Reaction was quenched with 5 mM EDTA and purified with Trizol. 100 ng demethylated small RNAs was used for library prep with a TGIRT Improved Modular Template-Switching RNA-seq Kit (InGex) following the manufacturer’s instructions. The included RNA/DNA heteroduplex for the template switching reaction by TGIRT contains a 3’ N-overhang on the DNA primer to promote reverse transcription of all small RNAs (as opposed to the T-overhang for targeting mature tRNAs described in (3)). PCR amplification was performed with Phusion polymerase (Thermo Fisher) with Illumina multiplex and barcoded primers synthesized by IDT for 12 cycles. Library was sequenced on a HiSeq4000. Data has been deposited at NCBI GEO, series GSE142393.

### DM-tRNA-seq bioinformatic analysis

tRNAscan-SE (61) was used to create the predicted tRNA gene library. Pre-tRNA sequences were defined by adding 50bp of genomic sequences on 5’ and 3’ ends of the predicted tRNA genes. Mature tRNAs were appended with a CCA tail and were clustered as described in (62). Illumina 150bp paired-end sequence data were subjected to quality control using BBDuk (https://jgi.doe.gov/data-and-tools/bbtools/). Sequencing adapters were removed from raw reads. Bases that have quality score lower than 30 were trimmed. Any reads that are at least 50 bp in length after quality control were aligned to mature tRNA reference using end-to-end mapping mode in Bowtie2 (63) (version 2.3.4.1). Reads that did not map to mature tRNAs were then mapped to the masked mm10 genome appended with pre-tRNA sequences. Two approaches were used for this mapping step. One is to use Bowtie2, with reads that map to multiple locations excluded from downstream analysis, followed by quantification of expression using SALMON (33). The other approach used SALMON to map the reads and quantify the expression levels, which includes the reads that map to multiple locations in estimating the expression levels. Raw counts were normalized using spike-in RNA species using R package RUVSeq (64). A generalized linear model was built to test for differential expression for each RNA species, using R package edgeR (65, 66).

### Chromatin immunoprecipitation

Cells from a 10 cm dish were crosslinked in 1% formaldehyde in PBS for 5 min at room temperature, quenched in 0.125 M glycine, and washed twice with PBS. Crosslinked cells were lysed with 1 ml ChIP lysis buffer (50 mM HEPES pH 7.9, 140 mM NaCl, 1 mM EDTA, 10% glycerol, 0.5% NP40, 0.25% Triton X-100) by rotation for 10 min. Nuclei were collected by centrifugation at 1700 x g for 5 min at 4°C, washed once with ChIP shearing buffer (50 mM Tris Cl, pH 7.5, 10 mM EDTA, 0.1% SDS), then resuspended in 1 ml of ChIP shearing buffer. Chromatin was sheared for 8 minutes using a Covaris S220 Focused-ultrasonicator at 140 power, 5% duty cycle, 200 bursts / cycle. Chromatin was spun at 15,000 x g for 5 min at 4°C and the pellet was discarded. 40 μg of chromatin was incubated with 10 μg rabbit polyclonal anti-POLR3A (Abcam ab96328) or rabbit IgG (Southern Biotech) overnight. 30 μl of mixed protein A and G dynabeads (Thermo Fisher) were added and tubes were rotated for 2 h at 4 °C. Beads were washed with low salt immune complex (20 mM Tris pH 8.0, 1% Triton X-100, 2 mM EDTA, 150 mM NaCl, 0.1% SDS), high salt immune complex (20 mM Tris pH 8.0, 1% Triton X-100, 2 mM EDTA, 500 mM NaCl, 0.1% SDS), lithium chloride immune complex (10 mM Tris pH 8.0, 0.25 M LiCl, 1% NP-40, 1% Deoxycholic acid, 1 mM EDTA), and Tris-EDTA for 5 min each at 4°C with rotation. DNA was eluted from the beads using 100 μl of elution buffer (150 mM NaCl, 50 μg/ml proteinase K) and incubated at 50°C for 2 hr, then 65°C overnight. DNA was purified using a Zymo Oligo Clean and Concentrator kit and used for qPCR analysis using the primers listed in Supplemental Table 1.

### Immunoblotting

25 μg of total protein was loaded onto with 4-15% Mini-PROTEAN TGX gels (Bio-Rad) and transferred to PVDF membrane. Blots were blocked with 5% milk/TBS+0.1% Tween-20 (TBST) and incubated with primary antibodies against Brf1 (Bethyl a301-228a), p53 (Cell Signaling 2524, clone 1C12), p-p53 (Cell Signaling 9284), or GAPDH (Abcam ab8245). Following incubation with primary antibodies, membranes were washed with TBST and incubated with goat anti-rabbit-HRP (1:5,000; Southern Biotech) or goat anti-mouse-HRP (1:5,000; Southern Biotech).

### Replicates and statistical analysis

This work contains cell culture-based assays, where biological replicates are defined as experiments performed on at least three distinct samples (cells maintained in different flasks). Technical replicates are defined as experiments performed on the same biological sample at least three times. The datapoint for each biological replicate performed is depicted in the Figures. Statistical analysis was performed using Prism 7 (Version 7.0c) software (GraphPad Software), and the exact test performed is described in the Figure legends. The criteria for including data involved the quality of the data for positive and negative controls.

## Acknowledgements

We thank Jie Li and the UC Davis Bioinformatics Core for performing the bioinformatics analysis, Michael Ly and Aaron Mendez for purification of AlkB enzyme, and Ella Hartenian for providing samples. We thank members of the Glaunsinger lab and Coscoy lab for discussion and reading of the manuscript. The model figure was designed using biorender.com. This research was supported by an American Cancer Society Postdoctoral Award 131370-PF-17-245-01-MPC to JMT and National Institutes of Health grants CA136367 and AI147183 to BG and GM120358 to IW. BG is an investigator of the Howard Hughes Medical Institute.

---

**Supplemental Table 1.**
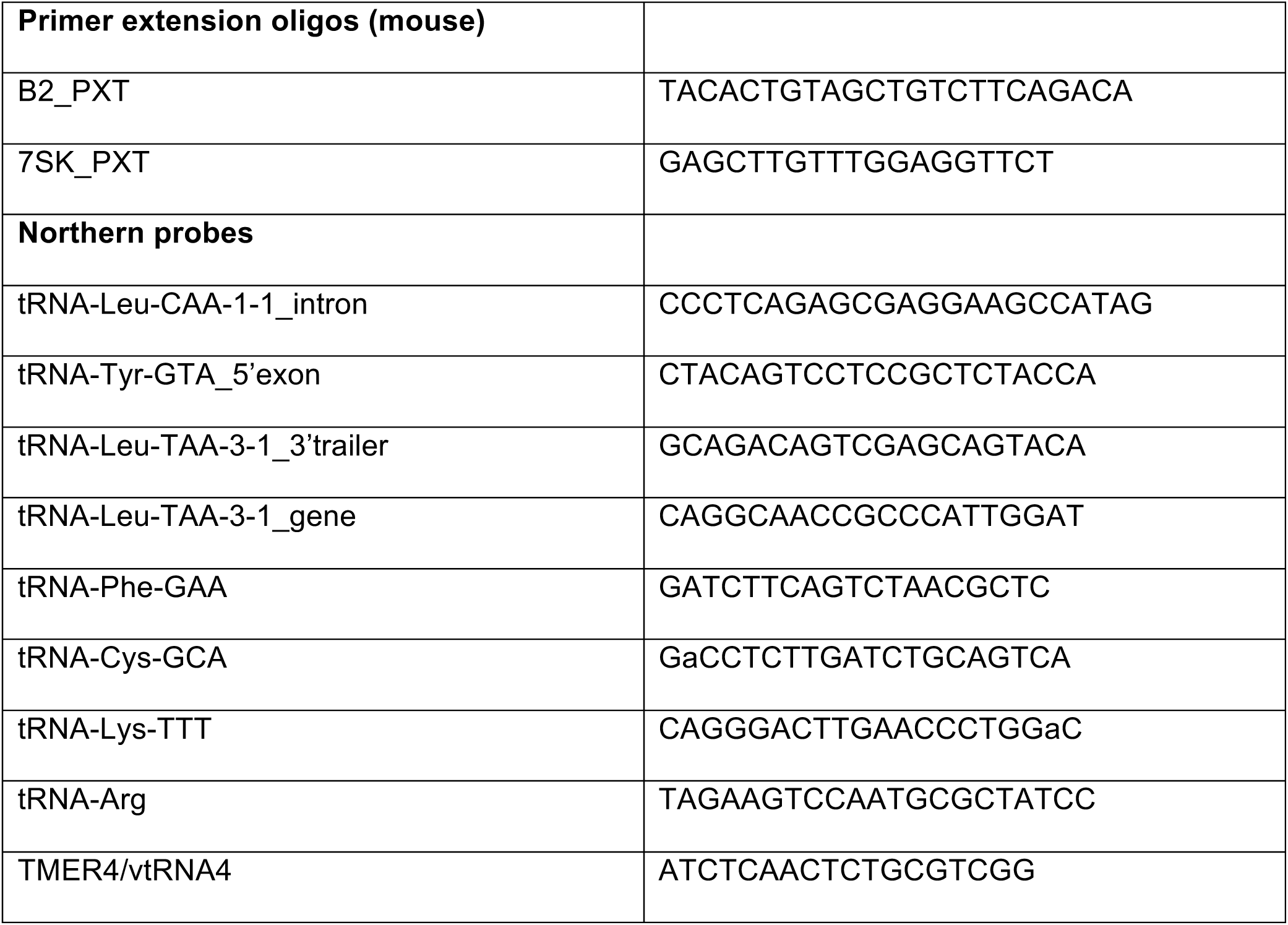
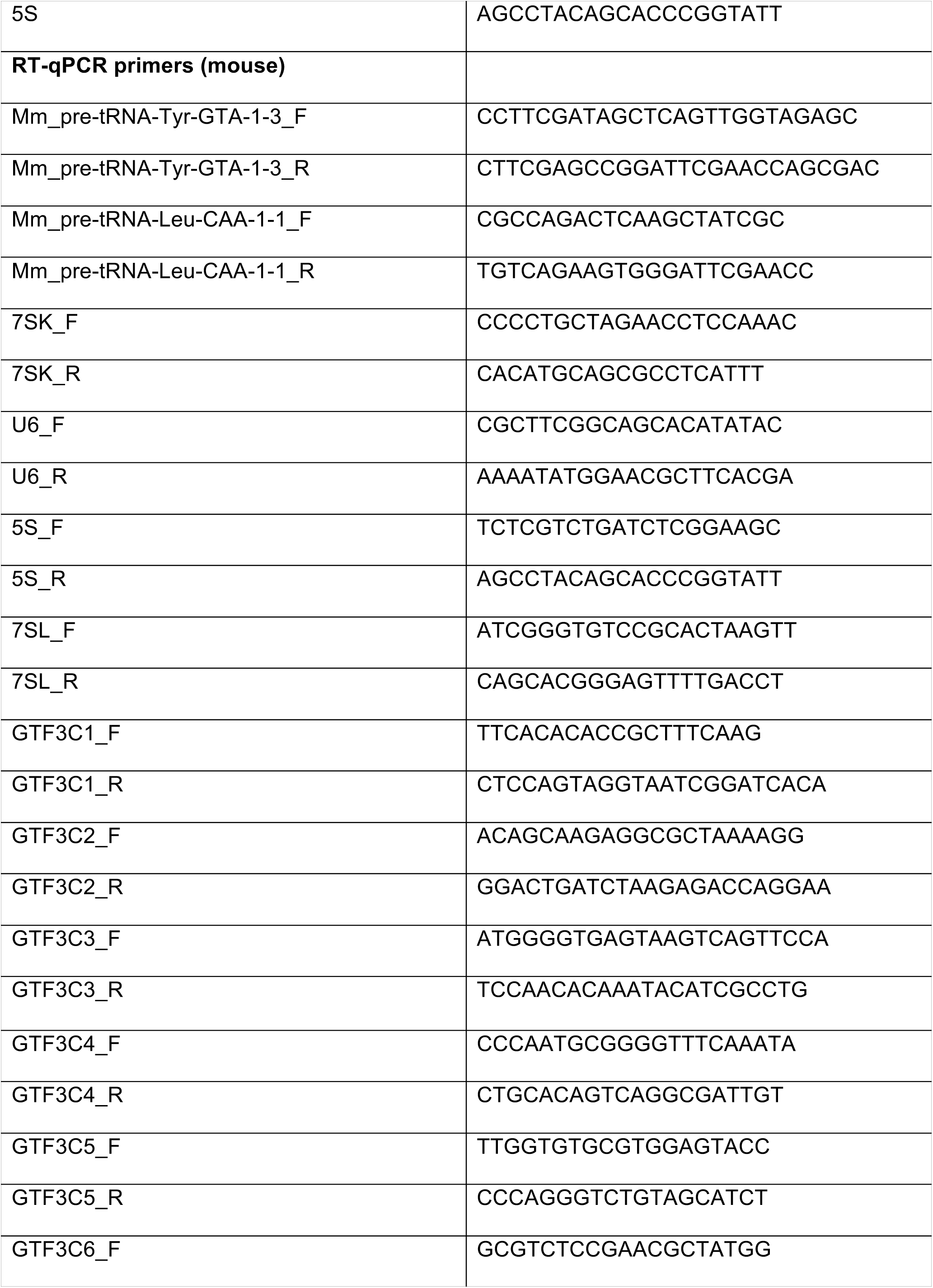
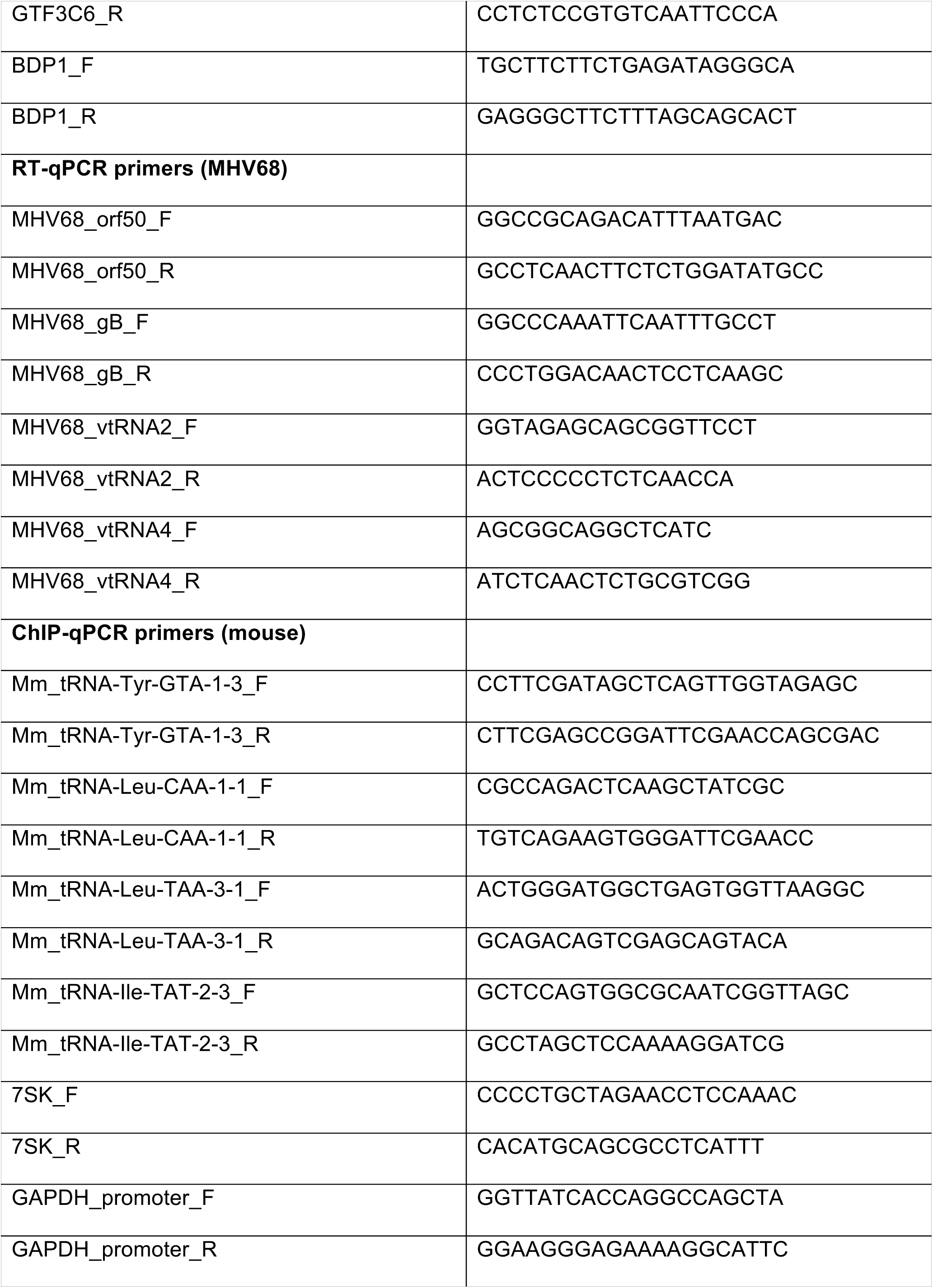
Oligos used in this study.

**Supplemental File 1.**
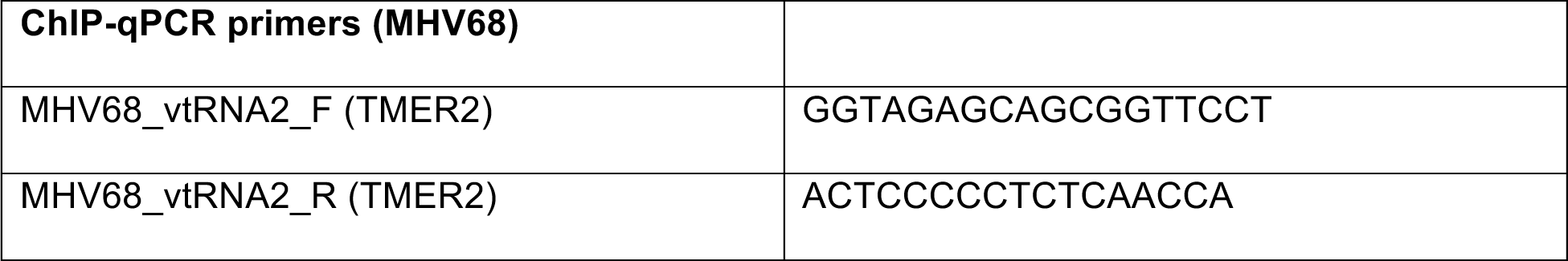
Differential gene expression analysis output from DM-tRNA-seq comparing MHV68-infected vs mock-infected MC57G fibroblasts.

**Supplemental Figure 1.**
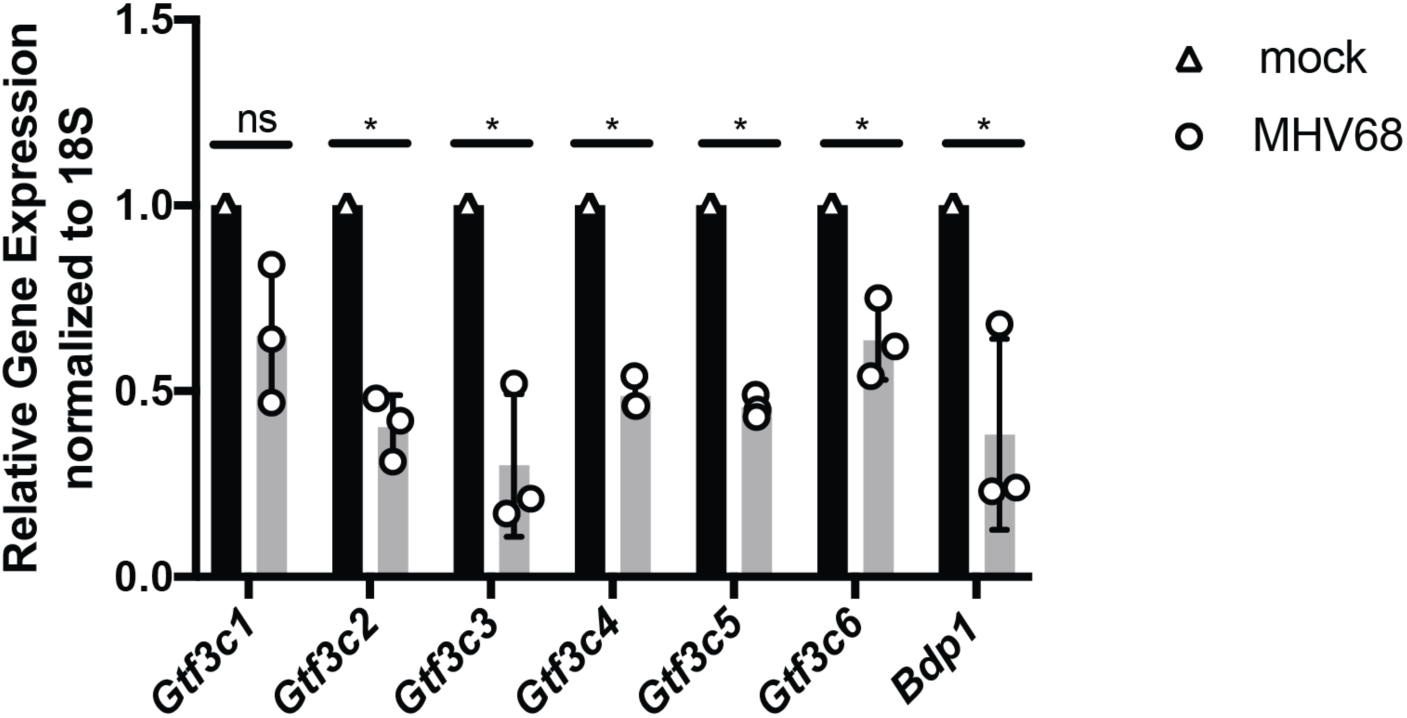
The levels of mRNAs encoding TFIIIB and TFIIIC subunits are decreased upon MHV68 infection. RT-qPCR was performed on MC57Gs infected at an MOI=5 for 24 h and were done in triplicate. Error bars show SD and statistics were calculated using an unpaired *t-* test on raw ΔCt values. *P<0.05.

**Supplemental Figure 2.**
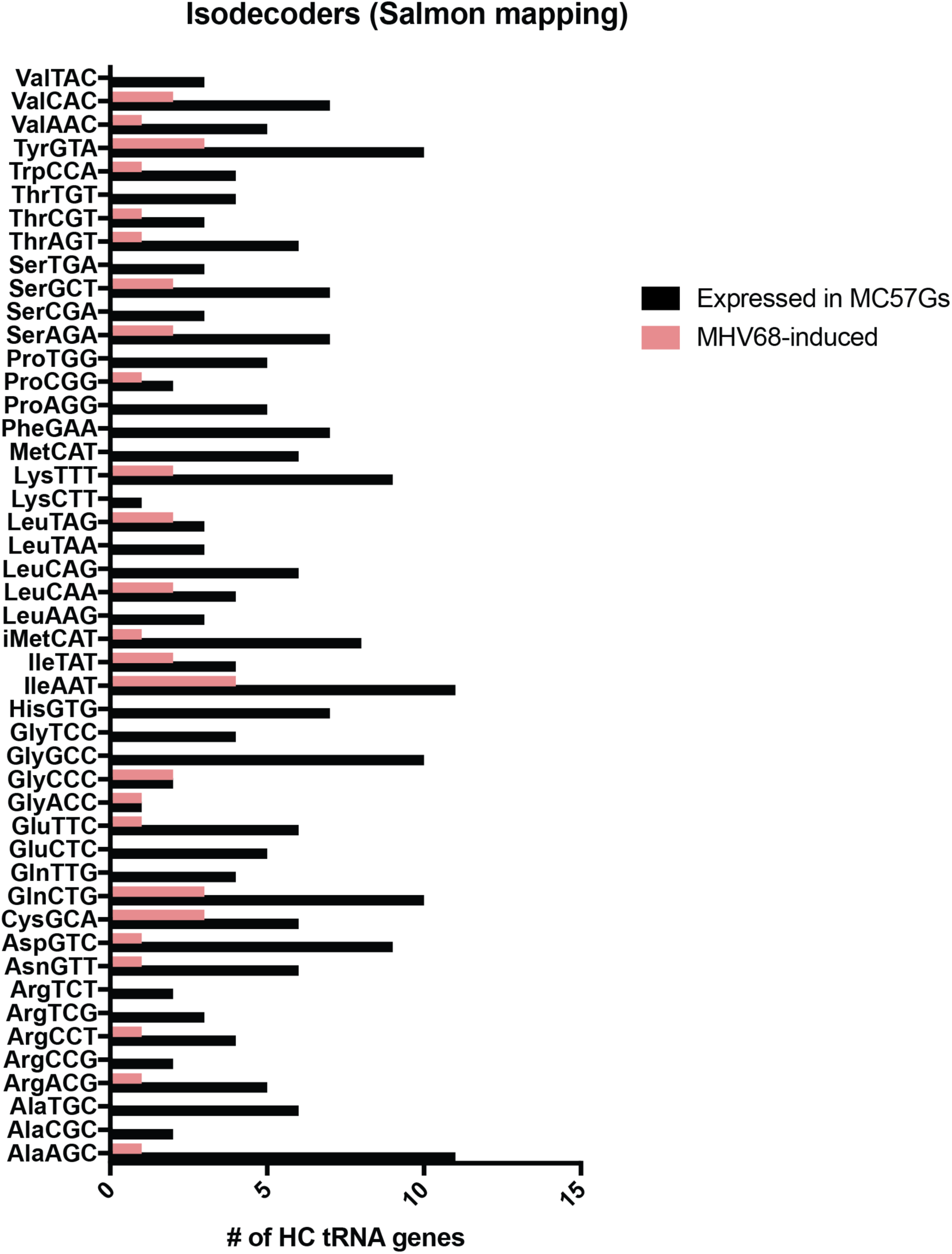
A variety of tRNA isotypes are upregulated at the pre-tRNA level in response to MHV68 infection. Pre-tRNAs detected by DM-tRNA-seq from each tRNA isodecoder are shown in black, while the number with increased pre-tRNA expression in response to MHV68 are shown in are pink. Most isotypes (tRNAs encoding the same amino acid), but not all isodecoders (tRNAs with same anticodon), are represented.

**Supplemental Figure 3.**
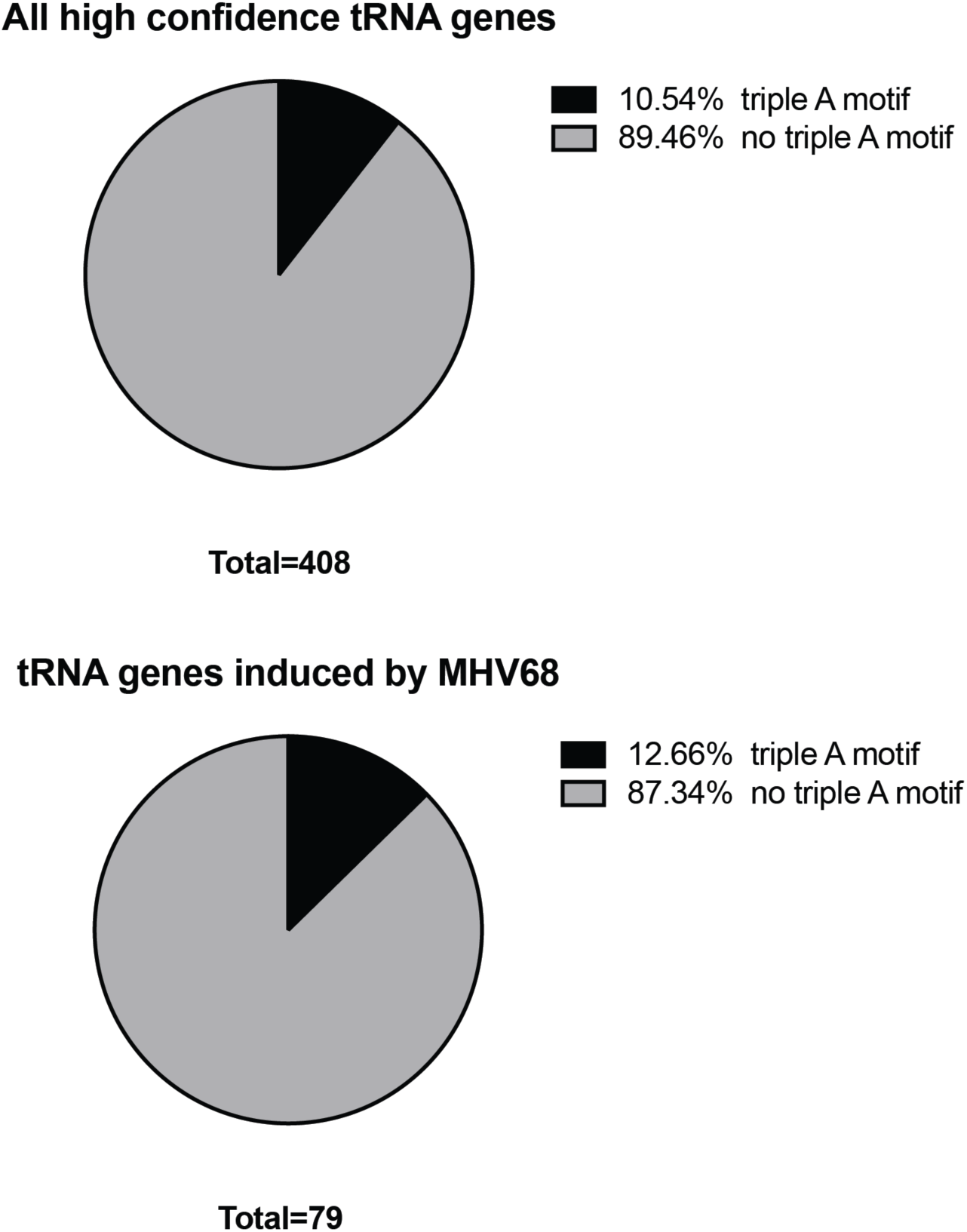
A triple A-box motif is not enriched in MHV68-induced tRNA genes. The triple A-box motif (TRGYNNARNTGGTRGARNAGNNG) identified in MHV68 TMERs is present in ∼11% of mouse high confidence tRNA genes as previously reported (35, 63). Both uniquely mapped and Salmon-identified MHV68-induced tRNA genes (n=79 genes) were analyzed for the triple A-box motif.

**Supplemental Figure 4.**
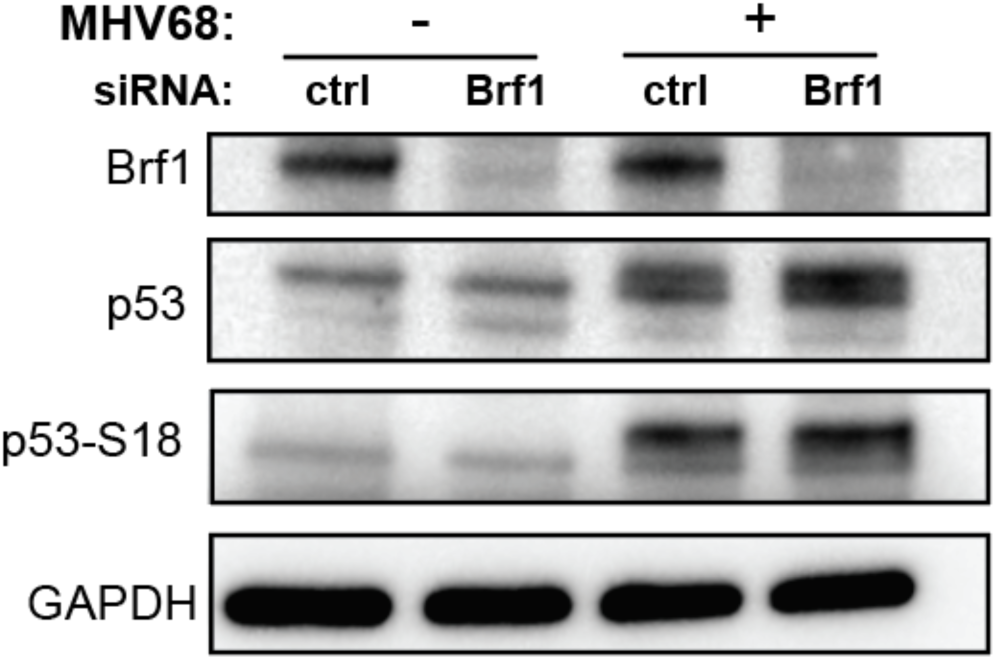
p53 phosphorylation at S18 during MHV68 infection is not dependent on Brf1 expression. NIH3T3 mouse fibroblasts were nucleofected twice with either control or Brf1-targeting siRNAs, then infected with MHV68 at an MOI=5 for 24h. Total protein was blotted for Brf1, p53, p53-S18, and GAPDH.

